# Neural Dynamics of Self-Control: Pre-Choice EEG Signals Predict Yielding to Temptation

**DOI:** 10.64898/2026.09.15.751887

**Authors:** Alisa Weng I Leong, Veit Stuphorn

**Author notes:** **Corresponding Author:** Veit Stuphorn.

## Abstract

Self-control is the capacity to resist self-defeating behavior in favor of long-term goals. Such behavioral control is fundamental to adaptive human behavior, yet its neural mechanism remains unclear. A key limitation of prior research is that tasks often conflate self-control with cost-benefit evaluation, making it difficult to isolate self-control encoding. We introduced a novel dynamometer-based task where participants chose between high-effort, larger-later (L) and low-effort, smaller-sooner (S) rewards. In some trials, referred to as temptation trials, participants could maintain their initial commitment (LL/SS) or switch (LS/SL). Twenty-six participants performed the task during a 32-channel EEG recording. Behavioral results showed a significant shift from L to S options in temptation trials, suggesting frequent yielding to temptation. EEG activity covaried with switching behavior. The late positive potential (LPP) differentiated stay (LL) from switch (LS) trials before initial choice, localizing to the posterior cingulate visual area and left somatosensory regions. Enhanced LPP in LS trials is thought to reflect heightened motivational conflict and attentional engagement with alternatives. Time-resolved multivariate decoding reliably discriminated LL from LS outcomes, peaking 304– 348ms post-option presentation. This was driven by distributed frontal and centroparietal networks, demonstrating that a neural bias toward yielding is established well before the initial choice at the network level. The EEG activity preceding switching predicted the final choice, reflecting reward anticipation rather than self-control signals. Together, these findings suggest that proactive neural states during deliberation, established before the choice, shape self-control success and failure.

**Significance statement:** This study demonstrates that self-control is a proactive process where neural states present long before a temptation must be overcome influence whether or not self-control is successful. Using a novel paradigm allowing preference shifts after commitment, we found that yielding to temptation is preceded by heightened late positive potential amplitude and distinct frontoparietal network patterns peaking at 304ms post-stimulus. Conversely, EEG activity preceding switching reflects reward anticipation rather than level of self-control. These findings suggest that self-control is a distributed neural mechanism that settles before the initial choice, providing new insights for studying deficits in addiction, gambling disorder, and obesity.

## Introduction

Self-control is the mental process of resisting immediate gratification to keep choices aligned with one’s long-term goals (Baumeister et al., 2018). Self-control plays a crucial role in decision-making by facilitating rational, goal-directed behavior through regulating impulses and considering long-term consequences (Baumeister, 2002; Boureau et al., 2015). However, the neural mechanisms underlying self-control are still unclear. Current tasks used to examine self-control suffer from the inability to disentangle self-control-related processing from other confounds, such as value-based evaluations and sustained attention throughout the experiment (Berkman et al., 2017). The delayed discounting task presents participants with a series of choices between a smaller reward available immediately and a larger reward contingent on a delay. However, choosing a smaller-sooner option over a larger-later one in this task does not inherently demonstrate a failure of self-control (Loewenstein & Carbone, 2024). Instead, it may reflect hyperbolic value discounting. In such cases, the participant is simply selecting the option with the highest perceived subjective value rather than failing to resist an impulse. Consequently, there remains a need for tasks that allow us to identify the influence of self-control independent of economic decision-making based on cost-benefit weighting.

Recent experiments in Rhesus Macaques used a novel paradigm, in which the animals first made a choice in a standard delayed discounting task, but occasionally had the opportunity to switch choices (Lee et al., 2024). Of interest were trials in which the animals initially chose an option that was more costly but optimal in the long run. However, when they were given the opportunity to switch, the sometimes maintained that choice and sometimes switched to a less costly, but suboptimal option. Under these conditions, maintenance of the long-term optimal choice indicates high self-control, while switching to the suboptimal low-cost choice indicates low self-control. Importantly, the initial choice was identical in both cases, so high and low self-control (maintenance or switching) was dissociated from value-based decision-making (initial choice). The supplementary eye field (SEF) encodes proactive self-control signals predicting success or failure of self-control in this task (Lee et al., 2024). Inspired by these findings, we adapted the paradigm into a novel human self-control task that allows us to compare trials in which participants maintained their commitment (high self-control) with trials in which they yielded to temptation (low self-control). We used electroencephalogram (EEG) recordings to characterize these dynamics in real time with high temporal resolution.

We focused on four primary questions: (1) Do early attentional processes differentiate successful from unsuccessful self-control? (2) Do motivational conflicts and attentional allocation during pre-choice predict self-control outcomes? (3) Can multivariate decoding of pre-choice EEG activity predict the eventual behavioral outcome above chance? (4) Do self-control-related neural signals persist beyond the pre-choice period into motor execution?

We utilized event-related potentials (ERPs) to investigate the neural dynamics, specifically the N1 and the late positive potential (LPP), both of which have been associated with successful self-control (Harris et al., 2013). The N1 (100-200ms) reflects early perceptual filtering (Ren et al., 2021). We hypothesized that N1 would differentiate successful self-control and unsuccessful self-control trials, with success associated with increased attention filtering of tempting attributes. Conversely, the LPP (400-800ms) indexes motivated attention and emotional significance (Hajcak et al., 2010). We hypothesized that unsuccessful self-control trials would exhibit an enhanced LPP amplitude, indicating greater engagement with both available options and more motivational conflict. Moving beyond focal effects, we employed multivariate pattern analysis (MVPA) to test whether outcomes could be predicted from the distributed neural activity

(Krönke et al., 2020). Finally, we examined the period preceding giving in to determine if these signals emerge at the time of giving in. By integrating these three complementary approaches, this study provides a comprehensive account of the neural dynamics of self-control as they unfold from initial deliberation to final action.

## Materials and Methods

### Participants

Thirty participants were recruited (12 males, 18 females; 18-23 years old with a mean age of 19.6) to perform the task. Written and informed consent was obtained from all participants, and all experimental procedures were approved by the Homewood Institutional Review Board of Johns Hopkins University. Participants were compensated with course credits. Two participants were excluded from both behavioral and EEG analyses due to failure to complete the experimental tasks. One participant was excluded from EEG analyses only due to the technical failure of the EEG recording files, and one participant did not have enough trials per condition for EEG analysis, resulting in 28 participants (12 males, 16 females; 18-21 years old with a mean age of 19.5) included in the behavioral analysis and 26 participants (12 males, 14 females; 18-21 years old with a mean age of 19.5) in the EEG analysis.

### Behavioral Task

The behavioral task was controlled using a computer equipped with MATLAB software and PsychToolbox-3 extensions (Brainard, 1997; Kleiner et al., 2007). The task was presented on a monitor that was approximately 57 cm away from the participants, and all the stimuli were limited to 5 degrees visual angle.

At the beginning of the experiment, participants were instructed to exert maximum force on a hand-clench dynamometer to determine their maximum voluntary contraction (MVC). Delays were defined as the duration for which participants must sustain 35% of their MVC on the dynamometer to obtain the reward for that choice. During the task (**Figure 1**), participants chose between a smaller-sooner (S) and a larger-later (L) option, with reward sizes corresponding to video durations of 1 s vs. 3 s and 3 s vs. 5 s. While the S-option delay was fixed at 500 ms, the L-option delays varied between 500, 900, 1300, 1700, and 2100 ms, appearing in 10%, 25%, 30%, 25%, and 10% of trials.

**Figure 1.**
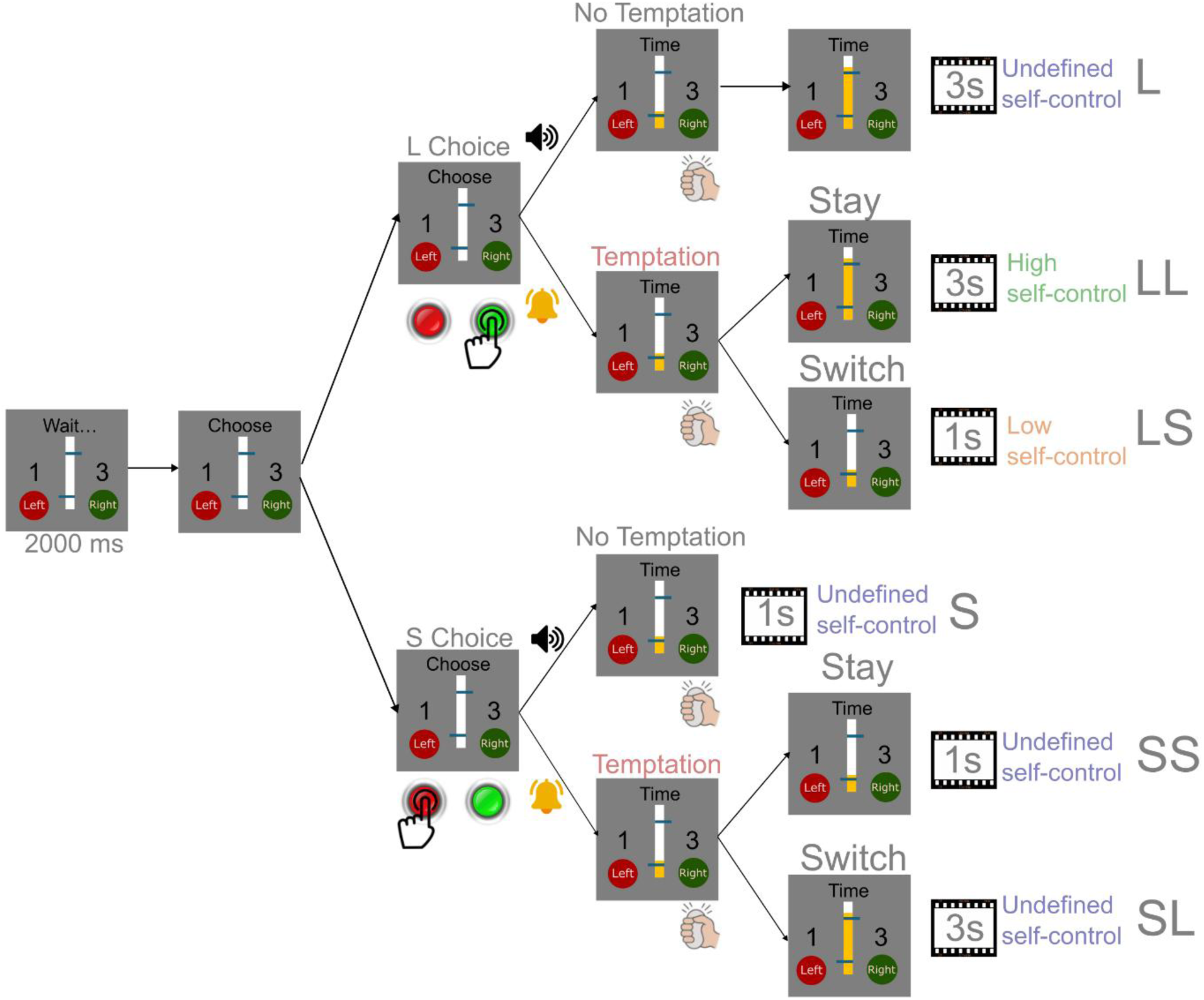
Self-control task design. After the options are presented for 2000 ms, the participants choose between the S and L options by pressing a button. The orange bar in the middle starts rising, when the subject starts pressing the dynamometer. It continually indicates how long the participant has already applied 35% of maximum voluntary contraction (MVC) on the dynamometer. The blue lines in the middle bar represent the delay for each option. They indicate the duration the participants must maintain a 35% MVC to earn the reward for the respective chosen option. In no-temptation trials, the participants have to stick with their initial choice and squeeze the dynamometer until the orange bar in the middle reaches the blue line of the chosen option. In temptation trials, the unchosen option remains available for participants. They can switch from their initial choice to the other one by squeezing longer or shorter on the dynamometer. Whether a trial is a no-temptation or temptation trial is indicated by an auditory cue after they make their initial choice.

Before the experiment began, participants selected videos that they would like to watch throughout the experiment as a reward. Video clips within each block were drawn consecutively from the same source video. The experiment consisted of 10 blocks, each with 50 trials: 25 temptation trials and 25 no-temptation trials, resulting in a total of 500 trials (250 temptation trials and 250 no-temptation trials in total).

Each trial began with the presentation of the options. Two horizontal lines on a central bar indicated the short and long delay (i.e., the costs). Numbers to the left and right indicated the video durations (the benefits). Above the bar, the text “Wait” was presented, and participants were instructed to think about their desired option during this period. After 2000 ms, the text changed to “Choose” and participants could start pressing the button to indicate their choice. When they pressed the button to indicate their choice, a sound was presented to indicate the type of trial (temptation or no-temptation). In a no-temptation trial, after making an initial selection, participants needed to apply at least 35% of the MVC of force on the dynamometer until the participant reached the delay of the associated option. If they failed to maintain 35% of MVC or release before they reached the delay of the associated option, the trial was deemed a failure, and nothing was shown on the monitor. The failed trial’s reward and delays pair was then repeated at randomly determined intervals throughout the remainder of the block to ensure the number of completed trials was consistent across subjects.

Temptation trials were presented randomly intermixed with no-temptation trials and were indicated with a different sound only after the participant made the initial choice. That way, the task setup was the same until the subject made the initial choice. Subjects could therefore not plan their initial choice based on whether a given trial was a temptation trial. Subjects were required to apply at least 35% of the MVC of force on the dynamometer to obtain the videos as well. The major difference in temptation trials was that participants could switch options (e.g., from L to S or S to L) by releasing the dynamometer earlier or maintaining the squeeze longer. If they switched from L to S (LS trial), the shorter movie would be played. If they switched from S to L (SL trial), the longer movie would be played. They could also choose not to switch their choice, which resulted in LL and SS trials. Maintaining the initial L option while the S option was available required self-control, as the S option represents an easier alternative with less cost. Trials in which subjects maintained their initial L choice, therefore, indicated high self-control, while trials in which the subject switched from the L to the S option (LS) indicate low self-control.

All the participants went through a brief training on the task to make sure that they could differentiate the tone of the no-temptation trial and the temptation trial. They were also trained to execute all types of trials (L, S, LL, SS, LS, and SL).

### Behavioral data analysis

The proportion of choosing the S option was calculated for each delay interval (500, 900, 1300, 1700, and 2100 ms) by dividing the number of trials in which the smaller reward option was selected by the total number of trials presented at that delay, separately for choice in no-temptation trials, initial choices in temptation trial (T1), and final choices in temptation trial (T2). To characterize the relationship between delay duration and choice behavior, we employed a logistic regression model to estimate the proportion of choosing the S option, p(S), as a function of the value of the L delay. The model was specified as:

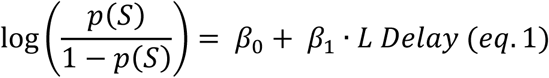

To assess statistical differences between choice in no-temptation trials, T1, and T2, we conducted pairwise comparisons using likelihood ratio tests on the fitted logistic models and complemented these with non-parametric binomial permutation tests (N = 1000 permutations). For the permutation test, we pooled the success counts and total trials from two conditions, then randomly reassigned these pooled trials to two groups while maintaining the original group sizes. The indifference point of the choice function was defined as the delay at which p(S) = 0.5, where the probability of choosing either option is equal.

### EEG data acquisition and preprocessing

EEG data were recorded in an electromagnetically shielded room. Thirty-two nonpolarizable Ag/AgCl electrodes were used from the Brain Products ActiCHamp system with actiCAP slim electrode caps, covering the whole scalp with uniform density using an electrode cap referenced to the Cz electrode during recording (ActiCHamp, Brain Products, Munich, Germany). Electrode impedances were kept below 10 kΩ. All EEG electrodes were recorded continuously in DC model at a 500 Hz sampling rate.

Data preprocessing was performed using the MNE-Python software (Gramfort et al., 2013) and the Autoreject library (Jas et al., 2017). Raw BrainVision EEG files were imported and digitized using the Standard 10-05 montage. EEG data were bandpass filtered between 0.1 and 40 Hz using a Butterworth IIR filter. To address transient sensor noise, the Random Sample Consensus (RANSAC) algorithm in the Autoreject library was employed to identify and mark bad channels based on spatial cross-correlations with neighboring sensors. A FIR high-pass filter at 1 Hz was applied to the data before the Independent Component Analysis (ICA). ICA was performed on continuous data using the Picard algorithm (Ablin et al., 2018). Components were automatically classified using the ICLabel neural network (Pion-Tonachini et al., 2019), and those identified as eye blinks or muscle artifacts were removed. After removing the artifact component, the data wes transformed back into sensor space. Marked bad channels were removed and interpolated. The data were re-referenced to the average and then sorted into epochs relative to target onset in two different time windows: (1) −200 to +2000 ms with baseline correction from −200 to 0 ms; (2) −700 to +200 ms with baseline correction from −700 to −500 ms. Peak-to-peak rejection thresholds were automatically estimated for each participant using the get_rejection_threshold function from the Autoreject package. Additionally, epochs with flat-line signals (activity within ±1 µV across the entire epoch) were excluded to remove periods of signal dropout or poor electrode contact.

### ERP analysis

For each participant, ERPs were computed by averaging epochs within each experimental condition. To address potential confounds arising from unequal trial counts between conditions (average number of trials: L: 127.3, S: 124.7, LL: 133.9, LS: 33.4, SS: 70.8, SL: 13.1), a bootstrap resampling procedure was used prior to comparison between conditions. For each participant, the condition with fewer trials was identified and set as the target sample size. Bootstrap resampling with replacement was then performed for 100 iterations from each condition, with the target number of trials drawn per iteration. The resulting bootstrapped averages were averaged across iterations to create a single, trial-count-matched evoked response for each condition and participant.

Statistical comparison between conditions was performed using nonparametric cluster-based permutation testing on a specific window of interest. For each participant, we first computed a difference wave by subtracting the two conditions. This approach controls the family-wise error rate (FWER) while maintaining sensitivity to effects that are physiologically plausible, specifically those that are clustered across adjacent electrodes and contiguous time points (Meyer et al., 2021). EEG data were sampled at 500 Hz, yielding a temporal resolution of 2 ms per sample. These individual difference maps were organized into a three-dimensional data matrix [Subjects x Times x 32 Channels] to serve as the input for the group-level analyses. A one-sample t-test against a zero baseline was then performed at each spatiotemporal point to identify sample-level differences. Adjacent sample points exceeding the statistical threshold (p < 0.05, two-tailed) were grouped into clusters based on spatial adjacency and temporal contiguity. The cluster-level statistics were defined as the sum of t-values within each cluster. We performed 1,000 permutations. In each iteration, the sign of the difference data for a random subset of participants was shuffled to construct a null distribution of the maximum cluster-level t-statistics.

### Source reconstruction

Source reconstruction was performed using sLORETA (Pascual-Marqui, 2002) in MNI space using the “fsaverage” template. A three-layer boundary element model (BEM) with conductivity values of 0.3, 0.006, and 0.3 S/m was constructed, and the source space was defined using octahedral subdivision (oct6, ∼4,098 sources per hemisphere). For each participant, electrode positions were co-registered to the “fsaverage” template using the standard 10-05 montage, and a forward solution was computed with a minimum source distance of 5.0 mm.

Noise covariance matrices were estimated from the baseline period using shrinkage and empirical methods. Inverse operators were constructed with loose orientation constraints (0.2) and depth weighting (0.8). To control unequal trial counts, bootstrap resampling, with 100 iterations, was applied, with both conditions subsampled to match the trial count of the smaller condition. For each bootstrap iteration, source activity was estimated using sLORETA (λ² = 1/9), and the resulting estimates were averaged across iterations to create trial-count-matched source estimates for each participant and condition. Grand average source activations were computed by averaging individual-level estimates across participants. The cortical region is defined by the Human Connectome Project multi-modal parcellation 1.0 (HCPMMP1) atlas (Glasser et al., 2016). To compare the difference between different conditions (i.e. L, S, LL, LS), we computed the within-subject difference between conditions for each parcel. To control family-wise error rate (FWER) across parcels, we performed a non-parametric sign-flip permutation test with 1000 permutations. The observed test statistic was a one-sample t-statistic computed for each parcel, testing the null hypothesis that the mean condition difference equals zero.

### Time-resolved decoding analysis

Time-resolved multivariate pattern analysis (MVPA; Grootswagers et al., 2017) was employed to investigate when and where brain activity patterns differentiate between high self-control conditions and low self-control conditions. For each subject, preprocessed EEG epochs were extracted. The data matrix had dimensions [Trials x Channels x Time points], channels = 32 electrodes, and time points corresponded to the epoch duration (-200 ms to 2000 ms at 500 Hz sampling rate). Binary labels were assigned to high and low self-control conditions (1 = LL, 0 = LS). To address the class imbalance, we used the Synthetic Minority Over-sampling Technique (SMOTE; Chawla et al., 2002). SMOTE was applied exclusively within the training partition of each cross-validation fold. We employed Linear Discriminant Analysis (LDA) with automatic shrinkage regularization for classification and evaluated using stratified 5-fold cross-validation.

For each subject and time point, we obtained Area Under Curve-Receiver Operating Characteristic (AUC-ROC) and balanced accuracy scores (one per fold), which were averaged to yield subject-level performance estimates. To assess the decoding accuracy significantly exceeding chance at the group level, we performed one-sample t-tests at each time point with the chance level. Chance level was set to 0.5, reflecting the binary nature of the decoder’s two possible outcomes. False discovery rate (FDR) was controlled over all time points at α = 0.05 using the Benjamini-Hochberg procedure. We also tested whether each electrode’s mean weight across subjects differed from zero using a one-sample t-test, with FDR correction for multiple comparisons across the electrodes. Positive weights indicate that higher activity at the electrode predicts the LL, while negative weights indicate prediction of LS. Weight magnitudes reflect the relative discriminative importance of each electrode.

## Results

### Effect of temptation on maintaining long-term goals

We estimated the probability to choose the S reward, p(S), as a function of the delay of the L reward (the cost), using logistic regression to determine the indifference delay, i.e. the delay at which the temporally discounted L reward is subjectively equal in value to the S reward. In temptation trials, the indifference point could be estimated for the initial choice (T1) and the final choice (T2). In no-temptation trials, the indifference delay was only determined for a single choice, comparable to the initial choice in temptation trials (T1). In the 1 s vs. 3 s movie clip trials (**Figure 2A**), the indifference point for choices in no-temptation trials was 1,610.3 ms and 1,711 ms for initial choices in temptation trials (T1). The difference between these conditions did not reach statistical significance (difference in indifference delay no-temptation-T1: -100.7 ms; 6% of initial indifference delay; permutation test, p = 0.062), indicating that participants did not know whether the current trial was a no-temptation or temptation trial when they made their initial decision.

**Figure 2.**
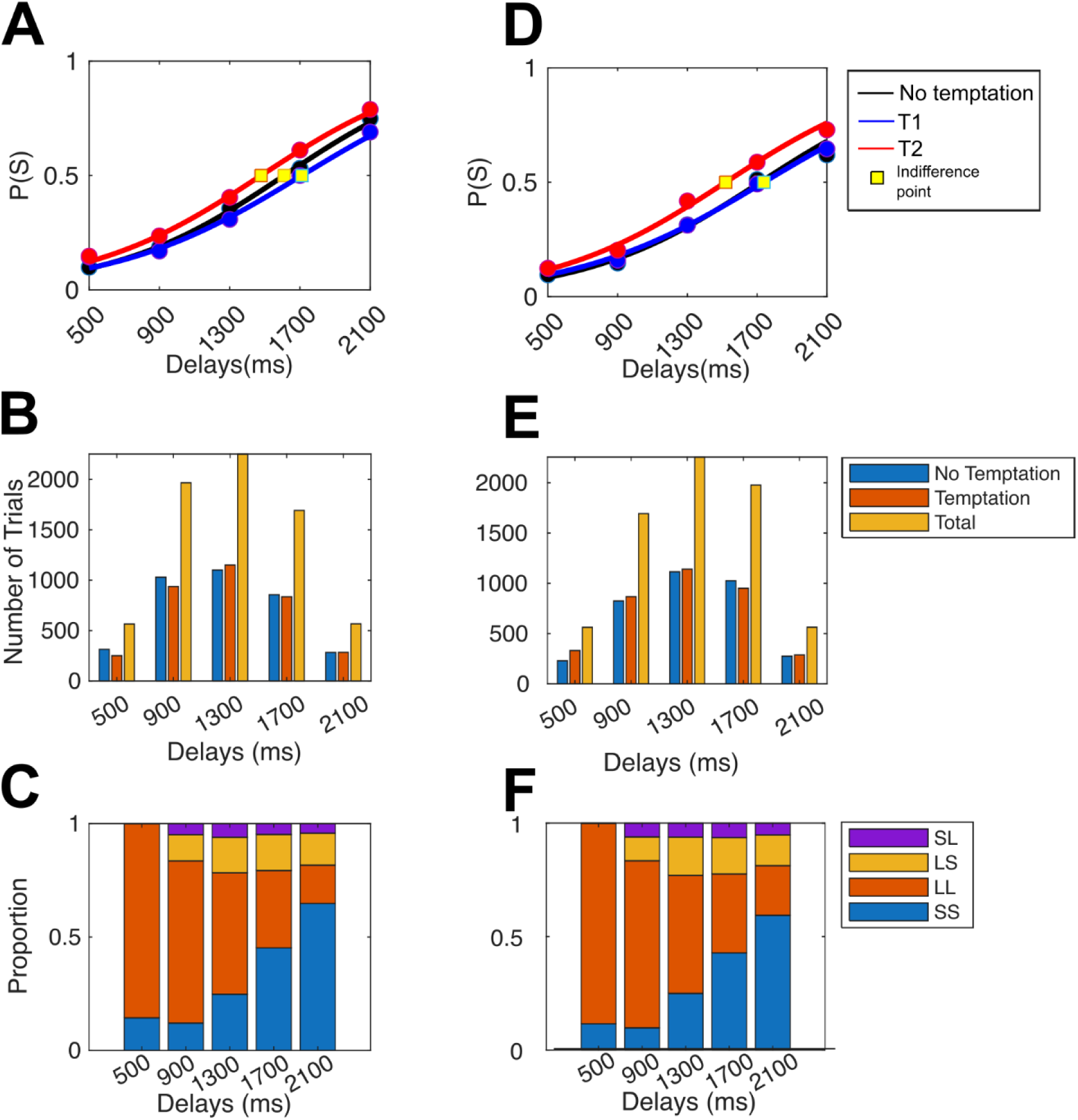
Behavioral results. (**A**) The proportion of trials where the S option was chosen as a function of L delays across all subjects in 1s vs. 3s trials. The black line is the choice function of the no temptation trial, the blue line (T1) is the choice function of the initial choice of the temptation trial, and the red line (T2) is the choice function of the function choice of the temptation trial. The yellow dot is the indifference point of the choice function. T1 and T2 are significantly different (permutation test, p < 0.001). (**B**) Distribution of the number of trials in different L delays across all subjects in 1s vs. 3s trials. (**C**) Proportion of trial types within temptation trials across all subjects for 1s vs. 3s trials. (**D**) The proportion of trials where the S option was chosen as a function of L delays across all subjects in 3s vs. 5s trials. T1 and T2 are significantly different (permutation test, p < 0.001). (**E**) Distribution of the number of trials in different L delays across all subjects in 3s vs. 5s trials. (**F**) Proportion of trial types within temptation trials across all subjects for 3s vs. 5s trials.

In temptation trials, the subjects are allowed to switch their preference. The difference between the indifference delays T1 and T2 reflected the effect of switches. Switches were rare (14.2% of all temptation trials). However, they caused a significant change in preference. The indifference delay for the final choice (T2, indifference delay = 1,478.8 ms) was significantly shorter than for the initial choice (difference in indifference delay T1-T2: 232.2 ms; 13.6% of initial indifference delay; permutation test, p < 0.001). This preference shift toward shorter delays reflects an asymmetry in the frequency with which the subjects switched between the reward types. There were substantially more switches from the larger-later to the smaller-sooner reward (LS; 73.3%) than switches in the opposite direction from the smaller-sooner to the larger-later reward (SL; 26.7%).

The same pattern can be found in 3s vs. 5s movie clip trials (**Figure 2B**). The indifference delay was 1,718.4 ms for no-temptation trials and 1,738.9 ms for initial choices in temptation trials (T1). T1 and no-temptation trial choice function overlapped, and statistical comparison revealed no significant difference (difference in indifference delay no-temptation-T1: -20.5 ms; 1% of initial indifference delay; permutation test, p = 0.446). In contrast, the indifference delay for the final choice (T2) of 1,519.5 ms was also significantly differed from T1 (difference in indifference delay T1-T2: 220.4 ms; 12% of initial indifference delay; permutation test, p < 0.001)

The shift in preference in the presence of temptation and the corresponding asymmetry in the direction of the shifts rules out error correction as the cause of the switches between the two options. Errors should be caused by random fluctuations in value estimation and should therefore lead equally often to incorrect choices of the S and L reward. The resulting error correction should produce equal rates of LS and SL switches, yielding a steeper choice function, but no shift in the indifference delay. Instead, the prevailing tendency to switch away from an initial L choice indicates that the L reward evokes conflicting motivation. On a longer-time horizon, the higher reward makes the L reward attractive, but on a short-time horizon, the higher cost makes the S reward more attractive. To maintain the choice of the L reward, self-control is required. The behavioral response to this temptation reveals therefore the level of self-control on that trial: persisting with the initial L choice despite this temptation (LL trials) indicates high self-control, whereas abandoning L for S (LS trials) indicates low self-control. The behavioral results in this human experiment are qualitatively similar to the results in the non-human primate experiment (Lee et al., 2024).

### Event-related potentials before choice reflect the level of self-control

A cluster-based permutation analysis was conducted to compare the EEG waveforms on LL and LS trials before the initial choice (**Figure 3**) between 400-1000 ms post-stimulus in the period between the onset of the options presentation and the “choose” signal. The analysis revealed a statistically significant cluster (p = 0.025) spanning from 692 to 796 ms post-stimulus onset, localized to posterior midline electrode sites (Oz and Pz, white dots on **Figure 3A**), consistent with the regions of interest of LPP.

**Figure 3.**
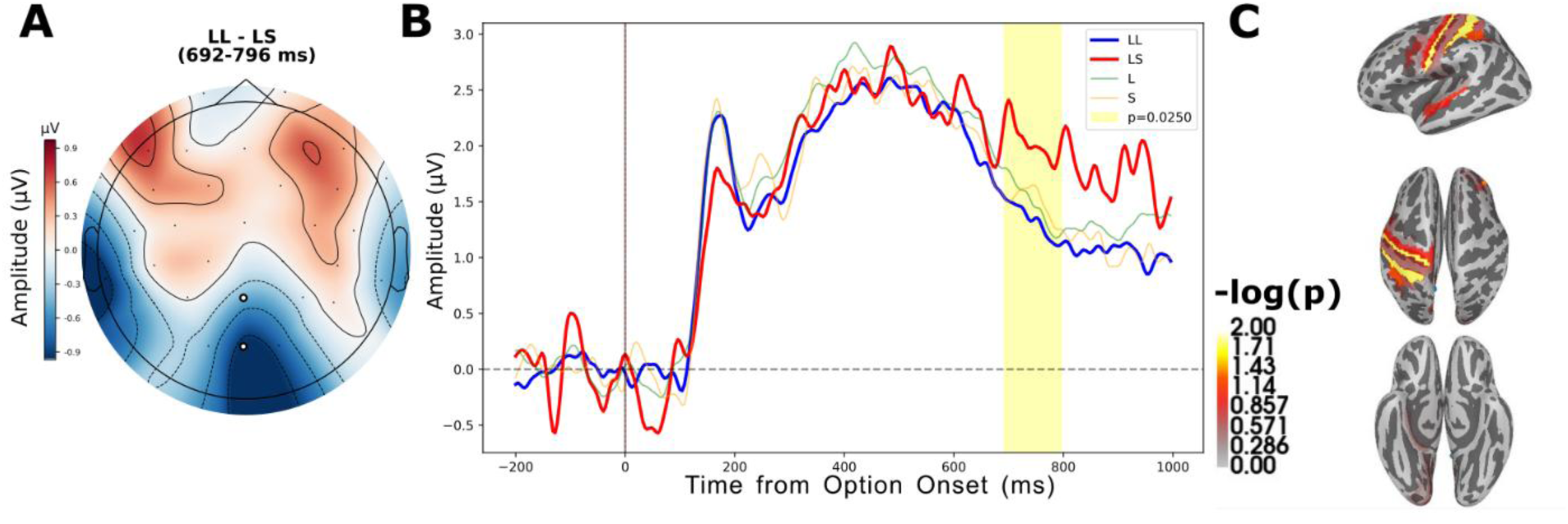
Cluster-based permutation results and source localization during the pre-choice period time-locked to option presentation. (**A**) Topographic map displaying the LL-LS amplitude difference averaged across the significant cluster duration (692–796 ms). White dots indicate electrode channels contributing to the significant cluster (p = 0.025). (**B**) Grand average EEG waveforms averaged across significant cluster channels for pre-choice conditions: LL (blue), LS (red), L (green), and S (yellow). The yellow shaded region denotes the temporal extent of the significant cluster. (**C**) Statistical parametric map (-log(p)) of sLORETA source localization for the LL-LS contrast during 692–796 ms. Source activity is overlaid on a standard brain template, revealing activation in the left posterior cingulate visual area, left area 3a, and left area 2 of the somatosensory cortex.

Grand average waveforms computed across the significant cluster channels (**Figure 3B**) revealed that the LS condition (red) exhibited a significantly more positive mean amplitude compared to the LL condition (blue) during the 692-796ms interval. Specifically, the LS condition demonstrated a mean amplitude of 2.3 µV (SD = 2.07 µV), whereas the LL condition showed a notably reduced positive deflection with a mean amplitude of 1.52 µV (SD = 1.53 µV). This pattern is indicative of an enhanced late positive potential (LPP) in the LS condition, potentially reflecting increased attentional engagement or evaluative processing when comparing options differing in both magnitude and delay dimensions. Importantly, this differential effect was specific to the LL versus LS comparison; no significant clusters emerged when examining the single-attribute L and S pre-choice conditions (green and yellow waveforms, respectively). Furthermore, the initial choice at the end of this period following the “chose” signal was identical (L). This difference could therefore not represent pre-motoric differences but instead reflected different levels of self-control.

Source localization analysis using sLORETA (**Figure 3C**) identified neural generators underlying the LL-LS difference within the 692-796 ms window defined by the HCPMMP1 atlas. The (-log(p-value)) statistical maps revealed significant source activity in the left posterior cingulate visual area (L_PCV; p = 0.010), as well as left somatosensory cortex regions including area 3a (L_3a; p = 0.015) and area 2 (L_2; p = 0.016), suggesting involvement of both posterior cingulate visual area and somatosensory areas in differentiating between these choice conditions during the pre-decision evaluation phase. Additionally, marginally significant activations were observed in the left anterior intraparietal area (L_AIP; p = 0.057) and left auditory area 5 (L_A5; p = 0.075), suggesting potential involvement of parietal regions associated with sensorimotor integration and decision-related processing.

To examine early attentional processing, the N1 amplitude was compared between LL and LS (time window 100-200 ms) over the occipital channels. No significant difference was observed between conditions (t (25) = 0.871, p = 0.392; **Figure 4**), indicating that the two trial types did not differ in early sensory or attentional processing of the choice options during this time window.

**Figure 4.**
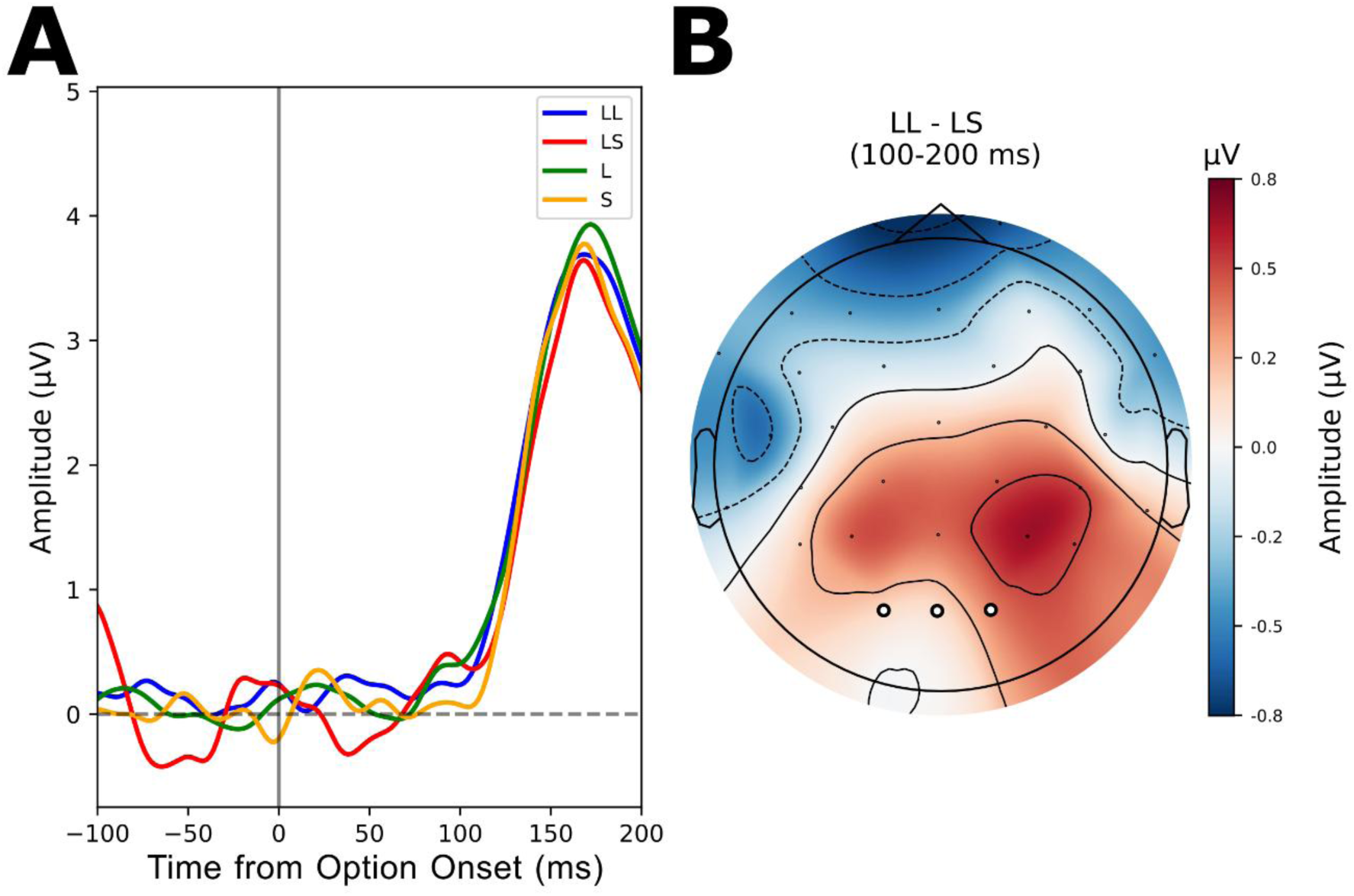
Cluster-based permutation on N1 window (100-200 ms). (**A**) Grand average EEG waveforms averaged across occipital channels for pre-choice conditions: LL (blue), LS (red), L (green), and S (yellow). (**B**) Topographic map displaying the LL-LS amplitude difference averaged across the N1 window (100-200 ms). White dots indicate occipital channels.

### Global EEG activity pattern before choice predicts outcome on temptation trials

To investigate whether self-control outcomes are reflected in global neural states, we applied time-resolved decoding across the full electrode array. This analysis revealed significant discrimination between LL and LS pre-choice conditions, demonstrating that distributed patterns of activity reliably predict behavioral outcomes associated with levels of self-control. Group-level analysis showed decoding accuracy significantly exceeded chance (AUC = 0.5) in a sustained time window (all p(FDR) < 0.05). Peak AUC occurred at 304 ms (mean peak AUC = 0.614, SD = 0.075, t (23) = 7.5, p(FDR) < 0.001, Cohen’s d = 1.531). Balanced accuracy showed a similar temporal profile (Mean peak accuracy = 0.587, SD = 0.087 at 348 ms, t (23) = 4.909, p(FDR) < 0.001, Cohen’s d = 1.002).

Analysis of LDA mean weights at the peak decoding window (304 ms) did not identify any electrodes with statistically significant contributions to classification after FDR correction, indicating that successful decoding was driven by a distributed neural pattern rather than activity at isolated electrode sites. Electrodes with the highest positive weights (indicating higher activity predicts LL choice) were primarily located over frontal and centro-parietal regions, including FC5 (mean weight = 0.24, Cohen’s d = 0.50), FP2 (mean weight = 0.24, Cohen’s d = 0.36), Cz (mean weight = 0.22, Cohen’s d = 0.27), P4 (mean weight = 0.21, Cohen’s d = 0.36), and CP5 (mean weight = 0.21, Cohen’s d = 0.43). Electrodes with negative weights (higher activity predicts LS choice) included Fz (mean weight = -0.28, Cohen’s d = -0.40), P3 (mean weight = - 0.25, Cohen’s d = -0.36), and P7 (mean weight = -0.17, Cohen’s d = -0.31), spanning frontal and left parietal areas. The topographic distribution of classifier weights revealed a broadly distributed pattern across frontal, central, and parietal electrode sites (**Figure 5C**), consistent with the involvement of widely distributed cortical networks in differentiating between LL and LS intertemporal choices rather than a single focal neural source.

**Figure 5.**
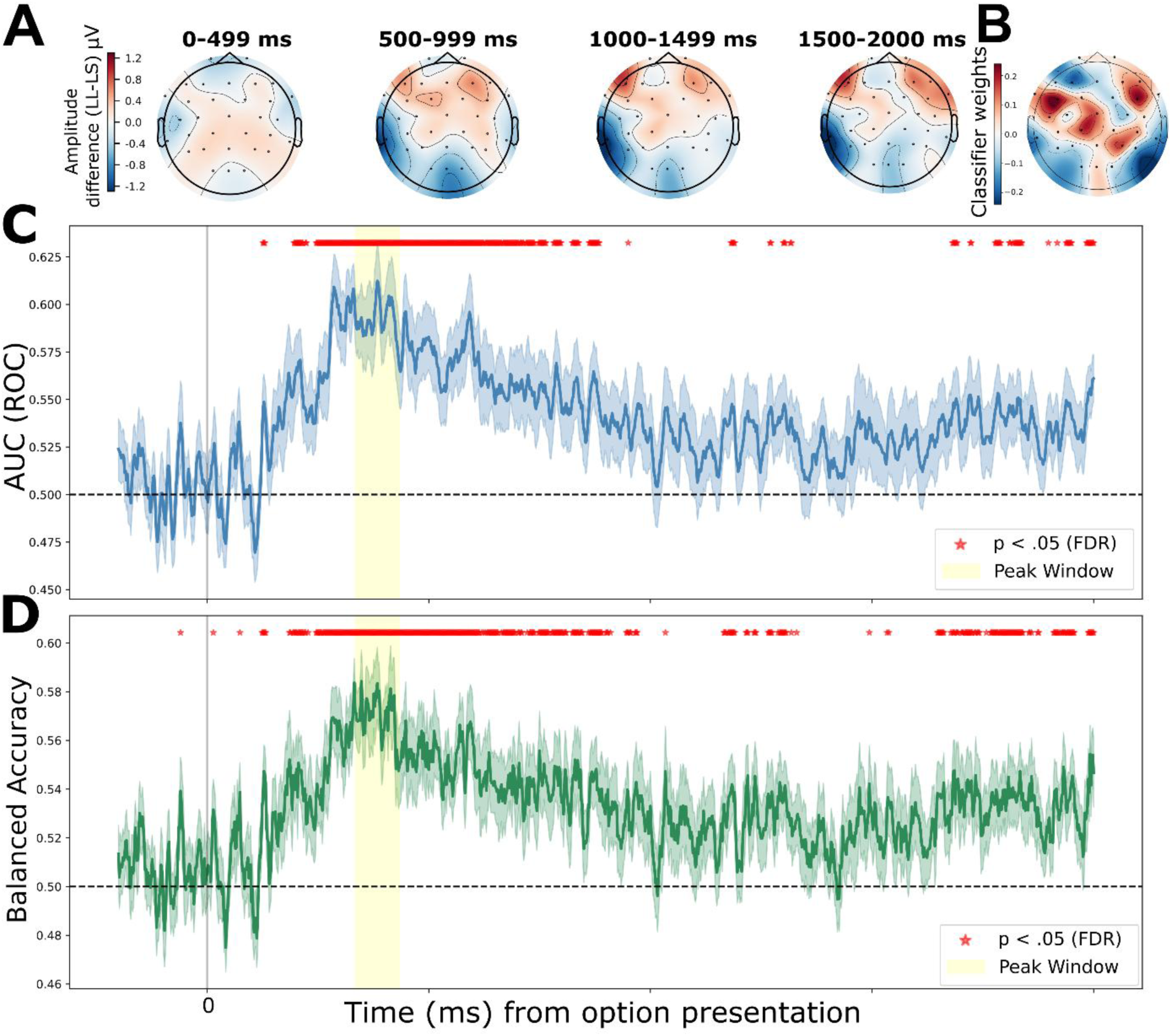
Decoding performance. (**A**) Representative topographic maps illustrating the average LL - LS amplitude difference across the pre-choice epoch, averaged in 500 ms windows. (**B**) Topographic distribution of classifier weights across all electrodes. Positive weights (red) indicate features predictive of LL choices, while negative weights (blue) indicate features predictive of LS choices. (**C**) Area under curve over the time course of pre-choice epoch with peak time point at 304 ms (Mean peak AUC = 0.614, SD = 0.075, t (23) = 7.5, p(FDR) < 0.001, Cohen’s d = 1.531). (**D**) Balanced accuracy over the time course of pre-choice epoch with peak time point at 348 ms (Mean peak accuracy = 0.587, SD = 0.087, t (23) = 4.909, p(FDR) < 0.001, Cohen’s d = 1.002). The stars represent the time points at which the p < 0.05 compared to the chance level.

### Reward value encoding during the late motor execution period

Spatio-temporal cluster-based permutation analyses were conducted to compare EEG activity between 4 conditions time-locked to the dynamometer release. When comparing the LL vs. LS conditions, the analysis identified two significant spatio-temporal clusters. The first spanned from -294 ms to 200 ms across 19 channels, and the second spanned from -288 ms to 200ms across 16 channels (**Figures 6A to D**). For the L vs. S comparison, three significant clusters emerged: Cluster 1 (p = 0.0010) spanned from -350 ms to 200 ms over 15 channels localized primarily around the parieto-temporal region; Cluster 2 (p = 0.0060) spanned from -380 ms to -142 ms over 14 channels in the frontal and frontocentral regions (**Figure 6E, F**); and Cluster 3 (p = 0.0050) spanned from -134 ms to 200 ms over 16 channels, also concentrated in the frontal region (**Figure 6G, H**). In the LL vs. S comparison, out of 9 total clusters, 2 were significant: Cluster 1 (p = 0.0010) spanned from -366 ms to 200 ms across 16 channels in the parieto-temporal region, while Cluster 2 (p = 0.0010) spanned from -348 ms to 200 ms across 16 channels in the frontal region. Finally, the LS vs. L comparison yielded 2 significant clusters out of 15 total. Cluster 1 (p = 0.0010) spanned from -262 ms to 200 ms across 17 channels in the frontal area, and Cluster 2 (p = 0.0010) spanned from -266 ms to 200 ms across 16 channels distributed across the central and parietal regions. Significant clusters consistently appeared across different condition comparisons, indicating that the magnitude or expectation of the final reward is the primary factor modulating the EEG waveforms during this timeframe. Thus, the observed EEG activity differences during the pre-release period seem to be primarily driven by the final reward (large or small reward) condition.

**Figure 6.**
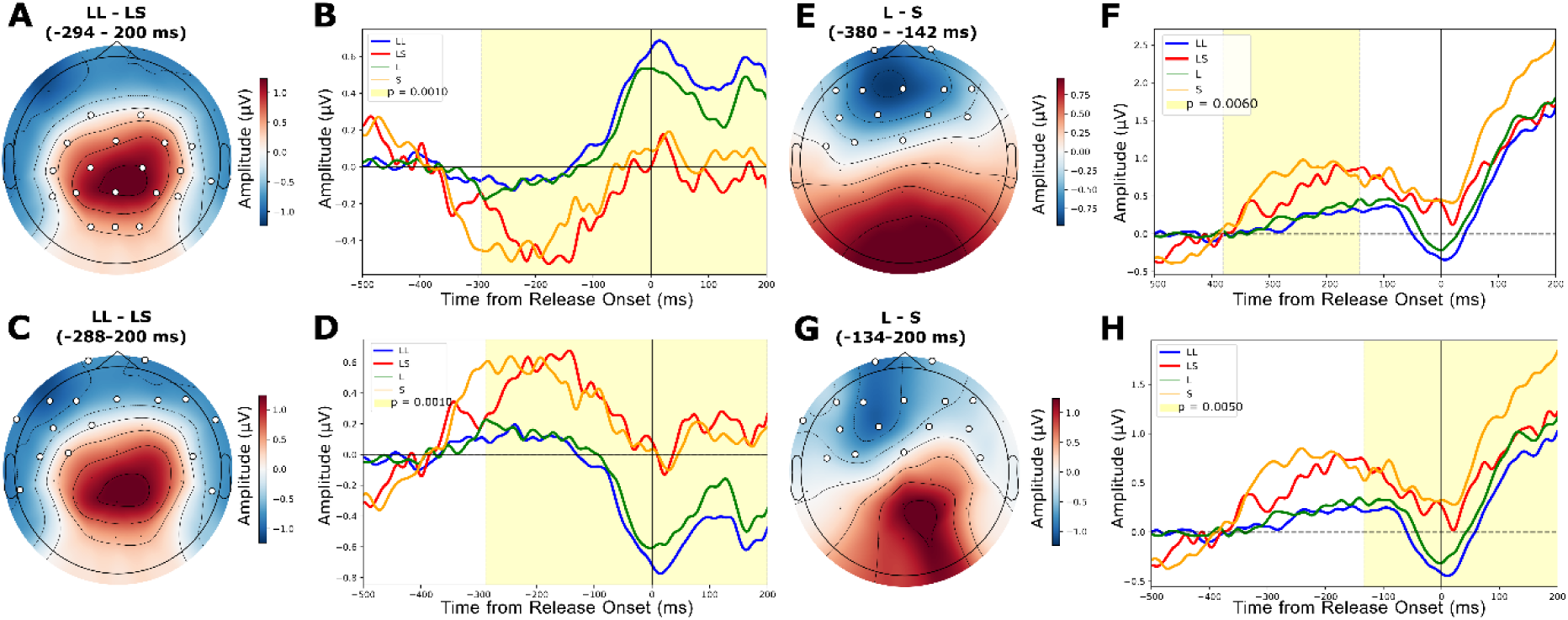
Spatio-temporal cluster-based permutation results and source localization of the pre-release period time-locked at the release time. (**A**) Topographic map displaying the LL-LS amplitude difference averaged across the significant cluster duration (-294-200 ms). White dots indicate electrode channels contributing to the significant cluster (p = 0.001). (**B**) Grand average EEG waveforms averaged across significant cluster channels for pre-release conditions: LL (blue), LS (red), L (green), and S (yellow). The yellow shaded region denotes the temporal extent of the significant cluster. (**C**) Topographic map displaying the LL-LS amplitude difference averaged across the significant cluster duration (-299-200 ms). White dots indicate electrode channels contributing to the significant cluster (p = 0.001). (**D**) Grand average EEG waveforms averaged across significant cluster channels for pre-release conditions. (**E**) Topographic map displaying the L-S amplitude difference averaged across the significant cluster duration (-380 - - 142ms). White dots indicate electrode channels contributing to the significant cluster (p = 0.006). (**F**) Grand average EEG waveforms averaged across significant cluster channels for pre-release conditions. (**G**) Topographic map displaying the L-S amplitude difference averaged across the significant cluster duration (-134-200 ms). White dots indicate electrode channels contributing to the significant cluster (p = 0.005). (**H**) Grand average EEG waveforms averaged across significant cluster channels for pre-release conditions.

## Discussion

This study demonstrates that self-control is a temporally extended neural process, with distinct signatures emerging well before the initial choice, and that failures of self-control are partly determined by the neural state during the preceding deliberation. This study uses a novel self-control task to investigate the neural dynamics of self-control by allowing shifts in preference after an initial commitment. Behaviorally, we observed a systematic shift of the final choice function in temptation trials towards less willingness to endure costs relative to the initial choice. This shift is caused by the fact that there were significantly more LS than SL switches. This asymmetry is critical, as it suggests that the shift represents a yielding to temptation rather than simple error correction. In support of these behavioral findings, our EEG results revealed that neural activity during the initial deliberation period, well before any switch occurred, significantly differentiated between successful (LL) and unsuccessful (LS) self-control outcomes. Specifically, the observed differences in late-stage evaluative processing (LPP) and global network-level patterns (MVPA) indicate that the seeds of self-control failure or success appear in the pre-choice period, that is, before the subjects knew that they could change their initial choice. This rules out any sensory or motoric processes as alternative explanations of these activity differences. Instead, they reflect proactive self-control levels.

The most significant local activity difference is the LPP differentiating LL and LS trials between 692 and 796ms post-option presentation in the pre-choice period, localized to posterior midline electrode sites (Oz, Pz), suggesting a difference in attentional engagement in the later stage of the pre-choice period. The LPP is a well-characterized ERP component indexing motivated attention allocated to emotionally or motivationally significant stimuli. Larger LPP amplitude reflects greater attentional engagement with salient stimuli and is enhanced when motivational conflict is present (Hajcak et al., 2010; Schupp et al., 2000). The heightened LPP for LS trials during the pre-choice period suggests that participants who ultimately yielded to temptation were experiencing greater motivational conflict and allocating more post-processing attentional resources to both options. This enhanced engagement with the choice alternatives, particularly the tempting smaller-sooner option, may have undermined subsequent self-control. Together, these findings suggest that self-control failure is not solely a failure of inhibition around the initial choice but is partly caused by the neural state of the participant during the preceding deliberation period.

To better understand the cognitive mechanisms driving this heightened attention, we employed sLORETA to identify the neural generators of the LPP effect. Source localization identified peak activity in the left posterior cingulate visual area (PCV), as well as left somatosensory cortex regions (S1), including areas 3a and 2. The involvement of the PCV is consistent with its established role in self-referential processing and integration of past experiences with current choices as part of the Default Mode Network (Brewer et al., 2013). In this task, PCV activation likely reflects that participants in the LS trial engage in a more vivid, self-focused recollection of the reward experience or effort experience, making them more vulnerable to temptation. Furthermore, the activity in the somatosensory cortex, specifically in area 2, supports the idea of motor imagery of the effort required in the task. Nierhaus et al. (2023) demonstrated that mental imagery of physical stimuli can trigger activation in the primary somatosensory cortex (S1). These results indicate that decreased self-control may be driven by a process of increased recollection of reward experience or effort experience.

In contrast to the LPP effects, the lack of significant N1 differentiation in our study suggests that early sensory filtering may be contingent upon specific task-timing properties. Our findings contrast with Harris et al. (2013), who reported enhanced N1 amplitude for high self-control individuals. They interpreted this as evidence for early attentional filtering of undesirable options. However, a critical procedural difference may account for this discrepancy. In Harris et al. (2013), participants could respond at any point during the option presentation window of up to 2.5 seconds. In the present paradigm, participants were required to view both options for a fixed 2-second period before responding, and each option comprised multiple attributes, effort level, and reward magnitude, which required concurrent evaluation. Given that response time has been shown to modulate N1 amplitude, with faster responses associated with larger N1 amplitudes (Schomaker, 2009), the fixed and uniform presentation duration in the present task likely resulted in comparable N1 amplitudes across LL and LS trials. Consequently, early attentional filtering may have been less functionally relevant in our task.

Complementing these component-specific findings, time-resolved MVPA analysis provided converging evidence that the neural state during the pre-choice period contains decodable information about the eventual behavioral outcome. Significant discrimination between LL and LS trials peaks as early as 304–348 ms post-stimulus onset, well before the initial choice, with higher positive weights (predicting LL) over frontal and centro-parietal regions and higher negative weights (predicting LS) over frontal and left parietal areas. The distributed nature of the LDA electrode weights suggests that the neural signature of self-control is not reducible to a discrete cortical source. Instead, this pattern implies that the information relevant to the decision is encoded across the collective activity of a broad ensemble of sensors.

Such a result indicates that self-control processes are likely maintained by the synchronized activity of a distributed network, where the interaction between regions is more predictive of behavior than the activation of any single node (Li et al., 2021). Previous research on self-control has identified multiple regions related to self-control, including the dorsolateral prefrontal cortex (dlPFC; Cosme et al., 2019; Hare et al., 2009; Harris et al., 2013), ventromedial prefrontal cortex (vmPFC; Moneta et al., 2023; Winecoff et al., 2013), inferior frontal gyrus (IFG; Han et al., 2018; Li et al., 2021), and dorsomedial prefrontal cortex (dmPFC; Jin et al., 2024) in functional magnetic resonance imaging (fMRI) studies. Many of these overlap with the Central Executive Network (CEN), suggesting that self-control emerges from coordinated activity across distributed prefrontal and parietal networks. This network-level organization is consistent with contemporary accounts of self-control as an emergent property of large-scale interactions between frontal and parietal regions rather than the product of a single cortical source (Brass & Haggard, 2007; Krönke et al., 2020).

The capacity of pre-choice activity to decode and predict the final decision parallels recent single-unit activity in the supplementary eye field (SEF) data reported by Lee et al. (2025), in which prefrontal activity during the pre-choice period reliably predicted the animal’s eventual decision to yield. Our results suggest that this proactive predictive signal is not species-specific but has a human EEG counterpart, extending the translational bridge between primate electrophysiology and human cognitive neuroscience.

Beyond the early decoding of choice, our findings reveal that the final stages of motor execution in the self-control task are characterized by anticipatory neural activity modulated by reward magnitude but not by self-control-related signals. Significant clusters were identified preceding the physical release of the dynamometer in LL compared to LS trials and L compared to S. Notably, the significance in the L vs. S comparison emerged earlier than in the LL vs. LS comparison, likely because the former lacked competing temptations, allowing for earlier reward-focused processing. This anticipatory activity closely mirrors the stimulus-preceding negativity (SPN), which is typically anteriorly distributed (Kotani et al., 2015), consistent with our observed cluster topographies (**Figures 6C–H**). Furthermore, these significant clusters persisted beyond physical release and bridged into the period of reward onset, suggesting that this activity is directly linked to reward processing.

Taken together, the present findings advance our understanding of self-control as a temporally extended neural process that unfolds well before the moment of overt choice, before temptation is present. Behaviorally, the asymmetric pattern of preference switches confirms that yielding to temptation reflects a motivational phenomenon rather than random variability or error correction. At the neural level, the heightened LPP for LS trials implicates increased motivational conflict and attentional engagement during deliberation as a precursor to self-control failure, while sLORETA localization points to the PCV as a key node integrating self-referential and reward-relevant information in this process. Complementing these ERP findings, time-resolved MVPA demonstrated that decodable neural signatures distinguishing successful from unsuccessful self-control emerge early, distributed across a frontoparietal network rather than any single region. Finally, pre-release ERP activity differentiating high reward and low reward trials suggests that reward anticipation is modulated by reward magnitude. Collectively, these results support a view of self-control as an emergent property of large-scale network dynamics. Furthermore, the results suggest that self-control is state-like and pre-determined during the pre-choice period. The results have the potential to assist populations characterized by self-control deficits, including those with substance use disorders, obesity, or attention-deficit/hyperactivity disorders, who may show altered or absent versions of the proactive neural signatures identified here (Balconi et al., 2024; Bullard et al., 2024; Fan & Jin, 2014). Applying the present paradigm in clinical samples could help determine whether self-control failures in these groups reflect impaired early proactive engagement, deficient conflict detection, or compromised motor-level inhibition.

## Acknowledgments

The authors would like to thank Dr. Erik Emeric for technical help with the experimental set-up, and Dr. Leyla Isik, Dr. Vikram Chib, Dr. Leo Chi U Seak, and Dr. Robert Ross for suggestions on analyses. This work is supported by NIMH grant R01MH137085 (VS), and an Albstein Research Scholarship (WIL).

